# How to improve foraging efficiency for restocking measures of juvenile Baltic sturgeon (*Acipenser oxyrinchus*)

**DOI:** 10.1101/483651

**Authors:** Maria Cámara-Ruiz, Carlos Espirito Santo, Joern Gessner, Sven Wuertz

## Abstract

Atlantic sturgeon (*Acipenser oxyrinchus*), also known as Baltic sturgeon, is considered missing or extinct in German waters. Current conservation efforts focus on re-stocking activities, but classical hatchery rearing may reduce the fitness of the respective juveniles. In this study, we evaluated if foraging efficiency can be improved by short term training. Over a period of 14 d, we kept individuals of the training group in a raceway and fed them chironomids buried in a small sand spot to stimulate benthic feeding behavior while fish of the control group were fed in tanks without substrate. Thereafter, each fish was transferred to a raceway entirely covered with sand. For feeding, chironomids were randomly buried in the sand. During the first 7 days, trained fish recovered the feed significantly faster than untrained fish of the control group. Gene expression revealed an up-regulation in *neurod1* in all brain regions after 14 d of training. Thus, this study suggests that foraging efficiency can be improved through short-time training thus improve fitness upon restocking into the wild.

## 1. Introduction

Sturgeons (Acipenseridae) were once native in all major rivers of the Northern Hemisphere, but over the last 100 years, they have shown a drastic decline due to fishing, habitat destruction and hydro-construction as well as pollution. In general, sturgeons exhibit an unusual combination of behavior and life history characteristics, particularly their late onset of maturity, making them highly vulnerable to anthropogenic impacts (Rochard et al., 1990). Nowadays, sturgeons are among the most endangered fish species worldwide (IUCN, 2018) and several restoration programs have been implemented. Baltic sturgeon (*Acipenser oxyrinchus)* has been indigenous to the Baltic region for the last 8000 years, but is now considered extinct in German waters. Restocking programs have been established as a part of the ongoing recovery efforts to reintroduce Baltic sturgeons into their natural habitats and, thereby, initiate self-sustaining populations (Gessner et al. 2011).

However, there are major concerns regarding early life experiences in artificial environments that may not produce fish prepared to face life in the wild (Johnsson et al., 2014). Studies have shown that classical hatchery rearing negatively affects fish fitness and eventually the performance and survival of the fish upon release into the wild (Sulak et al., 2014). It is known that in the natural environment, sturgeons experience stimuli that shape and influence brain plasticity, cognition and behavioral phenotype. In contrast, modern hatchery practices, including high stocking densities, predictable feeding regimes and uniform stimulus-poor rearing conditions, can result in impaired cognition and behavioural responses in sturgeons reared in captivity, leading to reduced fitness and survival (Ebbesson and Braithwaite, 2012).

In contrast to mammals, fishes display remarkable plasticity in brain neurogenesis, which remains active throughout adult life. As a consequence, fish are sensitive too and respond to changes in both social and environmental conditions (Ebbesson and Braithwaite, 2012). In their natural habitats, most fishes experience environmental challenges and are able to adapt their physiology and behavior in order to cope more effectively. Much of this flexibility is supported and influenced by cognition and neural plasticity. Furthermore, current literature on fish cognition indicates that many fish species are capable of learning and integrating multiple pieces of information that require more complex processes than associative learning (Ebbesson and Braithwaite, 2012).

Neuroscientists have payed special attention to the molecular mechanisms of neural plasticity associated with memory. This work has resulted in markers related to neural plasticity. Recent studies have indicated that proneural gene neurogenic differentiation 1 factor (*neurod1*) is a reliable measure of neurogenesis in fish and a useful indicator of the neural plastic changes associated with memory and learning (Rossi et al.,2006; Grassie et al., 2013). Moreover, brain-derived neurotrophic factor (*bdnf*) has an important role in neural plasticity through sculpting and refinement of synapses and through promoting neurogenesis and cell survival (Castrén and Rantamäki, 2009). Though not specific to the brain, proliferating cell nuclear antigen (*pcna*) is a marker for cell proliferation in the respective organ (Leung et al., 2005). Taking into account the functions of the genes previously mentioned (*neurod1*, *bdnf* and *pcna)*, they were of interest for this present study. In the wild, Baltic sturgeon is a benthic feeder which shows a digging behavior with help of the rostrum, preying on worms, shrimps and other invertebrates and making use of a powerful suction feeding mechanism (Carroll and Wainwright, 2003). Since most of their time is spent in waters with low visibility, feeding is performed by using a combination of olfactory, taste, tactile chemosensory cues and electroreceptors rather than vision (Mclean et al., 2013; Miller, 2004). The reduced importance of vision in feeding in sturgeons is supported by the observation that these fishes have relatively small eyes in relation to body size. In captivity, Baltic sturgeons are fed with artemia and thereafter with deep-frozen chironomids until they can be weaned on dry feed.

Taking into account that efficient foraging behavior is a key determinant of juvenile survival, the aim of this study was to determine whether natural foraging behaviour could be improved in hatchery-reared Baltic sturgeon following a short training period. In addition, this study also investigated whether this training resulted in positive changes in brain plasticity and cognition. To that end, following training, brains were sampled for gene expression analysis.

## 2. Materials and Methods

### 2.1. Experimental design

The experiments were performed at the experimental facilities at the Leibniz Institute of Freshwater Ecology and Inland Fisheries (IGB, Berlin, Germany) using fish from the sturgeon stock kept at the IGB. The experiment was conducted successively with 4 trained and 4 non-trained one year old Baltic sturgeon (*Acipenser oxyrinchus*) (TL 19-25 cm, 34 – 37 g) randomly distributed between 8 experimental raceway units (2.40 m * 0.225 m * 0.1 m) at a natural photoperiod and acclimatized for 7 days. Dissolved oxygen (8.41 – 9.16 mg/L), and temperature (18 – 20 °C) were measured daily, nitrit-nitrogen (0.002 mg/L) and total ammonia (TAN, 0.021 mg/L) every three days. Furthermore, 10 sturgeons were transferred to a constructed river stretch (6 m length, 1 m width, 0.2 m water depth, supplied by a Pontec Pondomax Eco 8000 pump) simulating close to natural conditions. The pond group was reared in an outside river stretch in order to compare the results of the trained and non-trained experimental groups to a more naturalistic environment (positive control).

Three experimental groups were established: non-trained, trained and pond (8 fish per group). The 8 raceways were assigned to non-trained and trained experimental group. The trained group had a sandy bottom (10 cm depth) while for the non-trained group the four raceways were left bare (10 cm depth). After a 7 day acclimatization period, during which the fish were fed chronomids, the training school started. The training was specifically designed to improve foraging behavior. Therefore, trained fish were fed chironomids hidden below a sand spot (< 10 cm), while the fish in the remaining bare raceways received chironomids on the bare tank bottom (non-trained). Before feeding in the respective raceway, fish were isolated by introducing a wall which was removed after feed had been introduced in the remaining part of the raceway. In the pond group, chironomids were hidden in the sandy substrate.

After 14 days, two sturgeon of each group were sampled for the gene expression, the remaining two fish were transferred to a raceway covered with sand (behavioral assessment).. For the assessment, chironomids were hidden in the sand and time until successful foraging was recorded for trained as well as for untrained fish (n=8). If chronomids were not successfully found within 120 min, food was removed. This assessment of foraging behaviour was repeated for seven consecutive days.

For the gene expression analysis, fish were euthanized with MS222 (300 ppm) followed by cutting through the spinal cord. Brains from 8 Baltic sturgeons per group (trained, non-trained, pond) were dissected and divided into three parts representing the three main brain regions (forebrain, midbrain and hindbrain). Samples were stored in RNA later at −80 °C for later gene expression analysis.

All experiments were in compliance with EU Directive 2010/63/EU and approved by the national authorities (G0305/15, Landesamt für Gesundheit und Soziales, Berlin, Germany).

### 2.2. Gene expression

Total RNA was extracted with TRIzol as described by (Reiser et al., 2011), including a DNase I digestion. Total RNA concentration and purity were determined in duplicates with a Nanodrop^®^ ND-1000 UV–Vis spectrophotometer. Purity was validated as the ratio of the absorbance at 260 and 280 nm (A260/280) ranging between 1.8 to 2.0. Moreover, integrity of the total RNA was checked by gel electrophoresis and, in 10% of all samples, on a RNA 6000 Nano chips with an Agilent 2100 Bioanalyzer. To eliminate potential DNA contamination, DNAse I digestion was performed in all samples prior to transcription. Next, mRNA was transcribed with MMLV Affinity reverse transcriptase (Agilent, 200 Units/μl) according to the manufacturer’s instruction. In 10% of the samples, the enzyme was substituted by pure H_2_0, serving as a control (-RT) to monitor DNA contamination.

Species-specific primers targeting elongation factor 1α (*ef1a*), brain-derived neurotrophic factor (*bdnf*), neurogenic differentiaton factor (*neurod1*) and proliferating cell nuclear antigen (*pcna*) were designed using the sequence information available. Specificity of the assays was confirmed by direct sequencing (SeqLab, Germany). Real-time PCR was carried out with Mx3005p qPCR Cycler (Stratagene), monitoring specificity by melting curve analysis. Full specifications of qPCR assays, including primer sequences are given in Table 1.

**Table 1.**
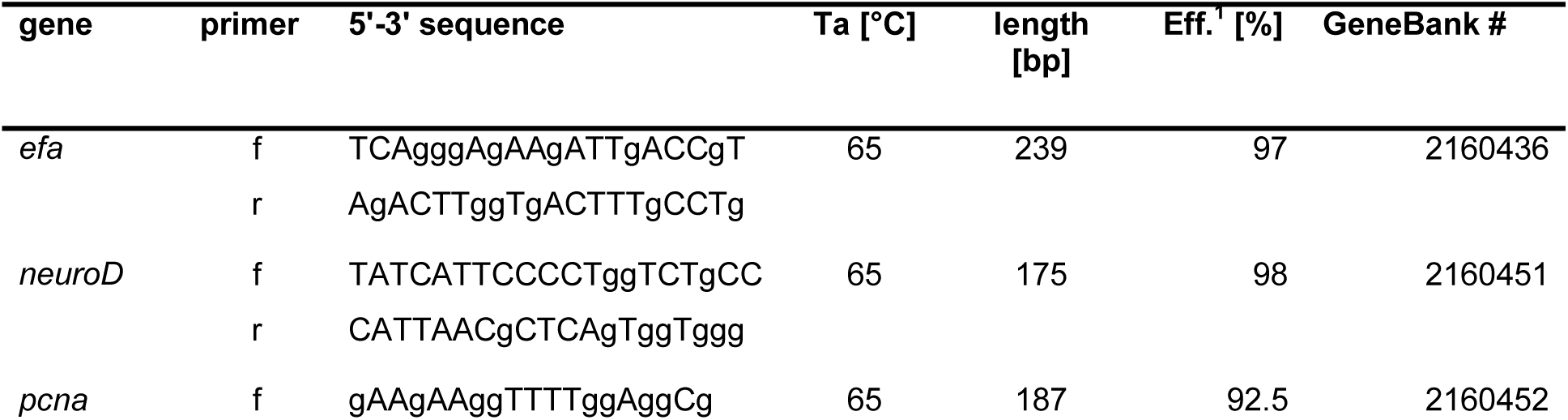
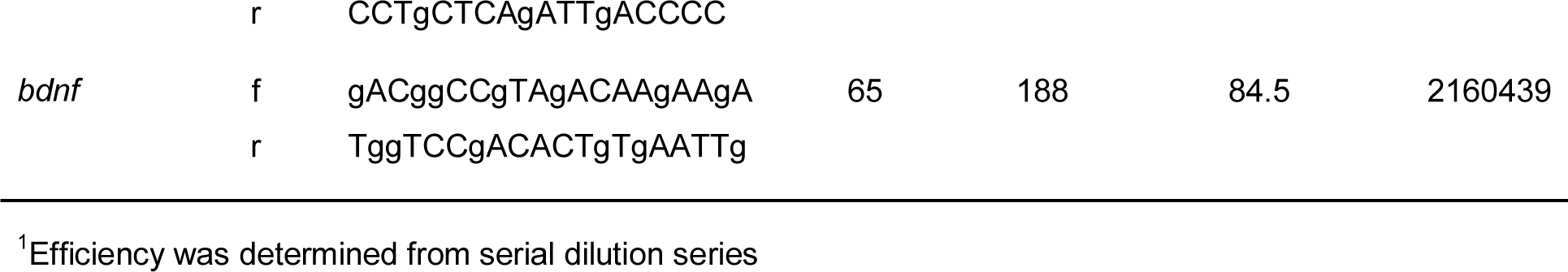
Specifications of qPCR assays including primer sequences, annealing temperature (Ta), amplicon length [bp], PCR efficiency (Eff) and NCBI accession number of the respective housekeeping (ef) and target genes: ef-elongation factor 1 a, neurod1-neurogenic differentiation factor, bdnf - brain-derived neurotrophic factor, pcna-proliferating cell nuclear antigen-

Briefly, 2 μL of the diluted sample (40 ng/μL) were used as template in 20 μL PCR mix [SYBR-Green I (Invitrogen), 200 μM of each dNTPs (Qbiogene), 3 mM MgCl 2 and 1 U Invitrogen Platinum Taq polymerase]. PCR conditions comprised an initial denaturation at 96°C for 3 min, followed by 40 cycles of denaturation at 96 °C for 30 s, primer annealing (for Ta, see Table 1) for 30 s and elongation at 72 °C for 30 s. PCR efficiencies were determined experimentally with a dilution series of a calibrator corresponding to 200 ng/μl. PCR assays for all individual samples were run in duplicate. Expression of target genes were calculated by the comparative CT method (ΔΔCT) according to (Pfaffl, 2001), correcting for the assay efficiencies and normalizing to elongation factor 1α (*ef1a*) as a housekeeping gene. Expression data are presented as fold increase of the respective control.

### 2.3. Data analysis and statistical methods

Data are presented as mean ± standard deviation (SD). Prior to statistical analyses, all data were tested for normality of distribution using the Kolmogorov-Smirnov/ Shapiro Wilk test and for homogeneity using Levene test. Data on the behavior were analyzed using T-test. The level of significance used was P ≤ 0.05. All statistical analyses were performed with GraphPrism statistical program.

## 3. Results

### 3.1. Behavioral assessment

In the behavioral assessment (Fig.1), significant differences were observed between non-trained and trained fish. While the time taken to initiate feeding successfully progressively decreased in both groups over the 7-day feeding study, this time was significantly less (p<0.05) in the trained compared to the non-trained fish at all time points (Fig. 1). On day 1, trained fish took 58±6 min to successful foraging whereas none of the non-trained fish recovered the chironomids within 120 min. After 7 d, chironomids were recovered after 8±2 min and 18±9min in trained and non-trained fish, respectively.

**Figure 1.**
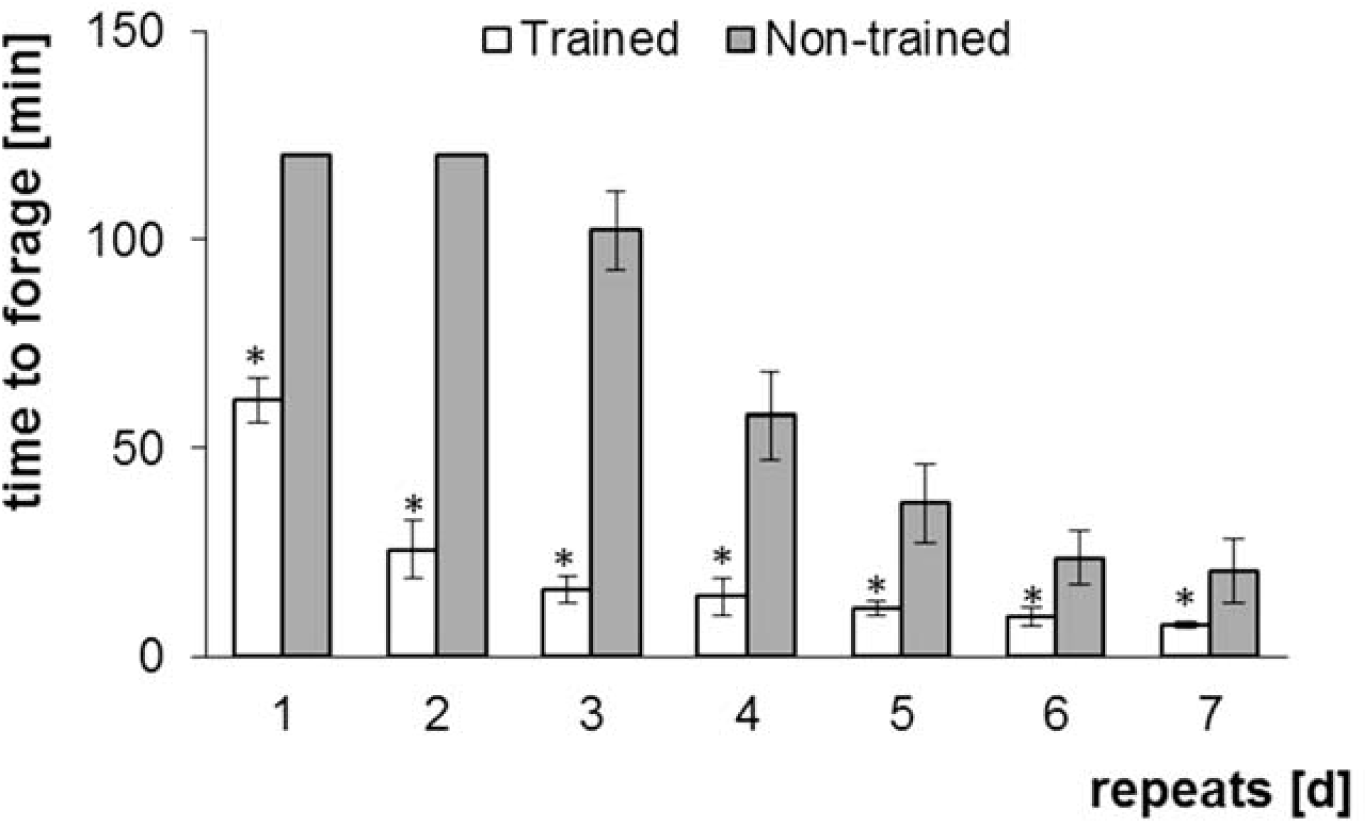
Behavioural assessment. Time taken by Baltic sturgeon (*A. oxyrinchus*) of the non-trained, trained and pond group to successfully forage. Significant differences are indicated by asterisk (Tukey’s, P < 0.05, n = 8)

### 3.2. Brain plasticity and cognition

Selected genes related to brain plasticity and cognition (*neurod1, bdnf, pcna*) were analyzed in all three brain areas of Baltic sturgeon. Regarding the forebrain region (Fig. 2), significant differences were observed in the expression of *neurod1*. In particular, there was an up-regulation in the trained and pond groups with respect to the non-trained group in Baltic sturgeon after 14 d of training (Fig 2A). Furthermore, *pcna* expression showed an up-regulation in the pond group in comparison to both trained and non-trained groups in the forebrain of Baltic sturgeon (Fig. 2B). Similar results were observed in both midbrain and hindbrain region (Figs. 3B & 4B) of Baltic sturgeon in which *neurod1* showed an up-regulation in both the trained and pond groups compared to the non-trained group after 14 d of training.

**Figure 2.**
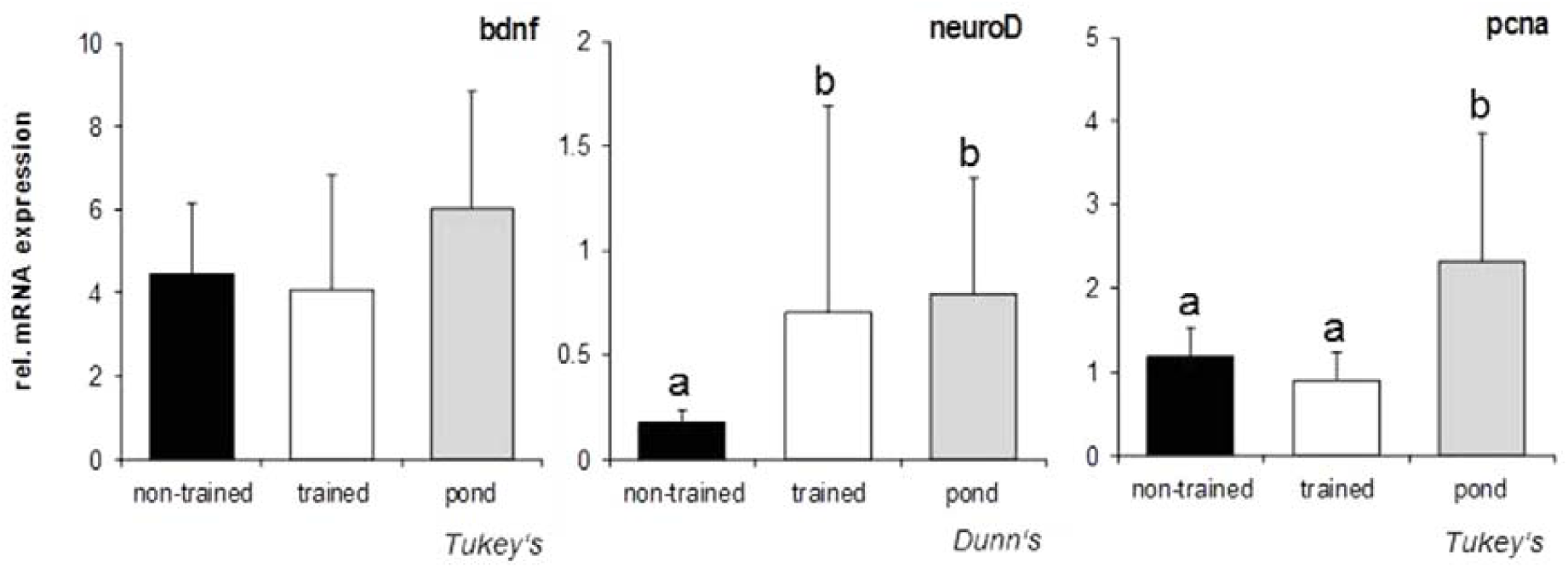
Gene expression, determined by qPCR, in the forebrain of Baltic sturgeon (*A. oxyrinchus*) of the non-trained, trained and pond group after 14 days of training. Data are expressed as fold relative. Groups with different subscripts are significantly different (P < 0.05, n = 8). Statistic test used (Dunn’s or Tukey’s) is indicated below the respective graph.

**Figure 3.**
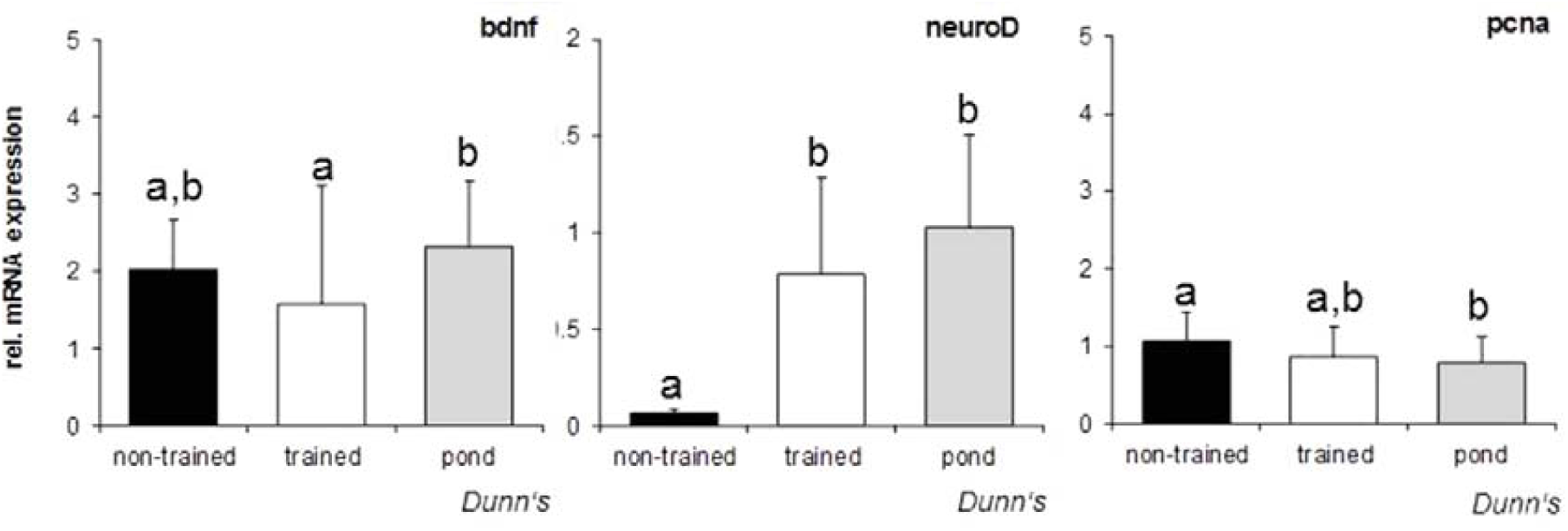
Gene expression, determined by qPCR, in the midbrain of Baltic sturgeon (*A. oxyrinchus*) of the non-trained, trained and pond group after 14 days of training. Data are expressed as fold relative. Groups with different subscripts are significantly different (P < 0.05, n = 8). Statistic test used (Dunn’s or Tukey’s) is indicated below the respective graph.

**Figure 4.**
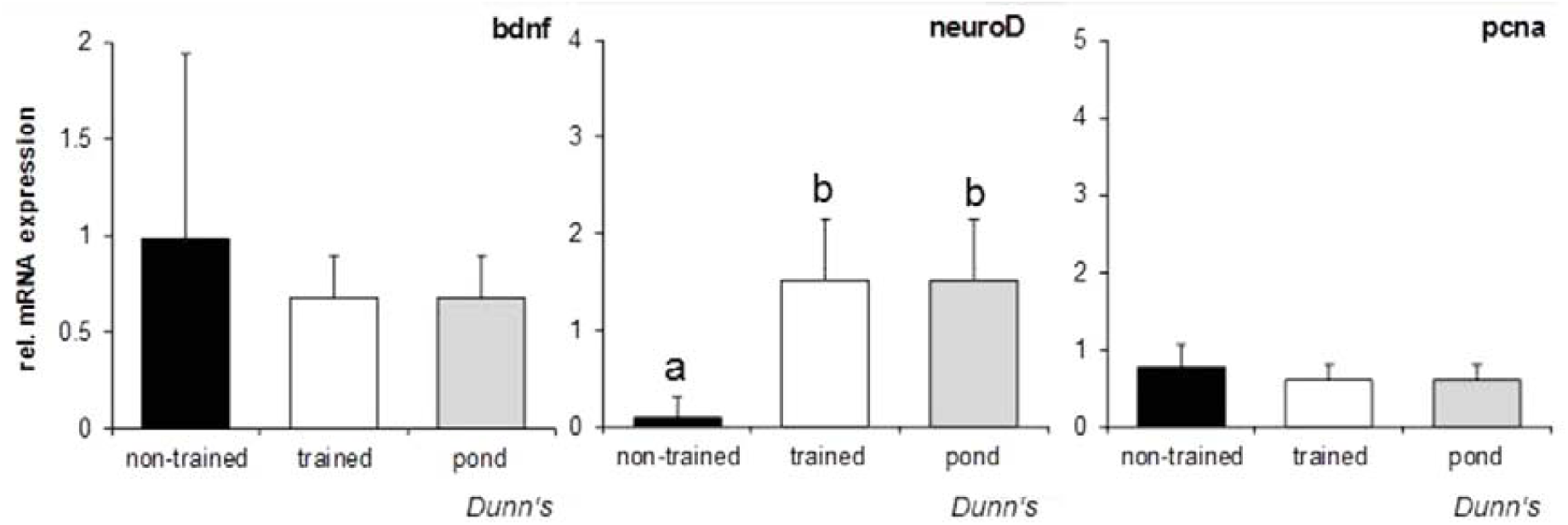
Gene expression, determined by qPCR, in the hindbrain of Baltic sturgeon (*A. oxyrinchus*) of the non-trained, trained and pond group after 14 days of training. Data are expressed as fold relative. Groups with different subscripts are significantly different (P < 0.05, n = 8). Statistic test used (Dunn’s or Tukey’s) is indicated below the respective graph.

## Discussion

Stocking still remains an important conservation tool to combat the continuing global decline in fish biodiversity (Pikitch et al., 2005). Hatcheries are a key element in the recovery plan for sturgeon and have been regarded as a temporary measure until more aggressive habitat restoration programs are established. In fact, hatcheries are currently the only viable option to increase sturgeon populations. Hatchery programs for sturgeon have demonstrated considerable success in collecting or developing brood-stock, spawning, and rearing juveniles. However, the success of sturgeon hatcheries for conservation will ultimately depend on how effectively the hatchery-reared sturgeon can adapt to the natural habitat following release (Brown and Day, 2002).

In general, when released directly into the rivers, the survival of hatchery reared juveniles from different fish species is only approximately 1-3% after a few months due to a combination of predation, starvation and other factors (Brown and Day, 2002; Chebanov et al., 2011). Furthermore, it is widely accepted that post-release survival rates of hatchery-reared fish are lower compared to their wild conspecifics (Campton et al., 1991; Svasand and Kristiansen, 1990). Fisheries scientists are increasingly convinced that the uniform stimulus-poor environment experienced during hatchery rearing is one of the main contributing factors for this reduction in fitness and post-release survival (Ellis et al.,1997; Masuda and Tsukamoto, 1998). Thus, research to improve the post-release survival of hatchery-reared juveniles through behavioral performance is needed in order to continue successful recovery plans.

Suggested methods for improving the survival of hatchery fishes include supplementary feeding with live foods, the provision of under-water feeders, sub-aquatic structure, natural substratum, etc (Maynard and Flagg, 1994) In the wild, Baltic sturgeon is a benthic feeder which shows a digging behavior in order to find worms, shrimps and other invertebrates from the substrate (Miller, 2004). Like all fish behavior, foraging relies on experience. The foraging skills of fishes become adjusted to ecological conditions through learning (Hughes et al., 1992; Warburton, 2003). Fish are able to learn to recognize prey, how to handle them and where they are likely to be located (Warburton, 2003). Results from the behavioral assessment of this study suggest that a short training period can improve the foraging ability of Baltic sturgeon by reducing the amount of time taken to successfully forage. This significant improvement was seen in the first 7 days, which can be critical when released into their natural habitat. This could be an important approach since when recaptured after release into the wild, hatchery fishes are often found to have empty stomachs (O’Grady, 1983; Johnsen and Ugedal, 1989).

The understanding of fish cognition and the role played by different brain regions has improved significantly in recent years. Fish brain remains plastic throughout their entire life and continues to be sensitive to both social and environmental changes. Most fishes experience challenges in their environmental and are able to adjust and adapt their physiology and behavior to help them cope more effectively. Much of this flexibility is supported by cognition and neural plasticity. Neural plasticity allows for the development and function of cognitive processes (Ebbesson and Braithwaite, 2012; Knudsen, 2004), and thus has a large role in the adaptation to changing environments. Current literature on fish cognition indicates that many fish species are capable of learning and integrating multiple pieces of information that require more complex processes than just associative learning (Ebbesson and Braithwaite, 2012). The most important brain area for these complex neural processes in fish is the dorsolateral telencephalon (Dl) (Durán et al., 2010; Rodríguez et al., 2002; Wullimann and Mueller, 2004), and has been recognized as the functional homologue of the mammalian hippocampus (Mueller et al., 2011; Mueller and Wullimann, 2009).

Studies using intermediate early genes (IEG) make possible to investigate which brain regions are activated during a particular cognitive process. In this study, three main genes were studied: neurogenic differentiation factor (*neurod1*), brain-derived neutrophic factor (*bdnf*) and proliferating cell nuclear antigen (*pcna*). Neurogenic differentiation factor (*neurod1*), is a member of a family of pro-neural genes, which is involved in the initiation and regulation of neural differentiation (Kiefer, 2005). Recent studies have shown that expression levels of *neurod1* mRNA is a reliable measure of neurogenesis in fish and a useful indicator of the neural plastic changes associated with memory and learning (Grassie et al., 2013; Salvanes et al., 2013). Brain-derived neurotrophic factor (*bdnf*) is the most abundantly expressed member of the nerve growth factor family, neurotrophins, and has an important role in neural plasticity through sculpting and refinement of synapses and through promoting neurogenesis and cell survival (Castrén and Rantamäki, 2009). It has recently been shown that environmental challenges alter *bdnf* expression in the telencephalon of Atlantic salmon (Vindas et al., 2017). Regarding proliferating cell nuclear antigen (*pcna*), it can be used as a marker for cell proliferation (Leung et al., 2005).

In this study, *neurod1* was up-regulated in all brain regions in both the trained-group and fish raised in the semi-natural pond, in comparison to the non-trained fish. This might be an indication of the stimulation of cognitive processes such as learning and memory as a result of the training method. Thus, it indicates that the fish from the trained group generally learnt to locate the prey. These results are in agreement with the results found in the behavioral assessment.

Currently, the main limitation of stocking programs for most fish species is the high level of post-release mortality. The most critical period seems to be the immediate days following release as hatchery-reared fish generally display reduce life fitness traits such as foraging and anti-predator behaviour.. In our study, it was demonstrated that a short time period of training could potentially help Baltic sturgeon in their process of learning to successfully forage, which could be a first approach to improve restocking practices. Since rearing conditions are highly important in stocking for conservation, hatcheries should aim to produce juveniles that are morphologically, genetically, behaviorally and physiologically similar to the stock they pretend to enhance and recover. Furthermore, restoration programs require a variety of information on sturgeon and thus, it is interesting to produce and keep up to date extensive reviews of the literature. Further work is needed in order to determine the survival of sturgeon reared under alternative hatchery-rearing practices taking into account other key factors in sturgeon survival.

## Conclusion

To our knowledge, this is the first study that looks into foraging training in Baltic sturgeon to improve fitness for re-stocking purposes. We observed that both behavioural and physiological parameters were improved by a short-term training period. This improvement could significantly help Baltic sturgeon survive in the wild, since the highest percentage of mortality happens during the first days post-release.

## Acknowledgments

The authors thank Eva Kreuz for her contribution in the laboratory. The study was supported by the European Training Network of the Marie Sklodowska-Curie Actions ITN “Improved production strategies of endangered freshwater species”. This project has received funding from the European Union’s Horizon 2020 research and innovation program under the Marie Sklodowska-Curie grant No [642893]

## Author contributions

The experiment was conducted by C.E.S. The laboratory analysis was carried out by M.C.R. M.C.R. wrote the first draft of the manuscript. S.V. supervised the project. The manuscript was revised by all co-authors.

## Table of Abbreviations

d-: days
TL-: total length

